# NADH as a cancer medicine

**DOI:** 10.1101/019307

**Authors:** Michael D. Forrest

## Abstract

We propose that NADH will exert a specific kill action against some cancers. NADH is a natural metabolite. We envisage a low side effect profile and that NADH therapy will, additionally, combat the wastage and weakness of cancer patients, which can be the cause of death in some cases. Significantly, NADH can be administered orally and has already cleared clinical trials, all be it for other pathologies.

**Background:** Aerobic respiration consists of glycolysis in the cytoplasm and in the mitochondria: the Krebs cycle and oxidative phosphorylation (OXPHOS) [1, 2]. It requires O_2_ and net yields 30 ATPs from one glucose molecule. Anaerobic respiration, consisting of glycolysis only, does not require O_2_ but produces merely 2 ATPs from one glucose molecule. When O_2_ is available, normal animal cells tend to favour aerobic respiration because of its higher ATP yield.

---

Aerobic glycolysis is the sole use of glycolysis to produce ATP, even in the presence of O_2_. Pyruvate and NADH, the outputs of glycolysis, are not fed into the further, mitochondrial steps of the Krebs cycle and OXPHOS to gain a higher ATP yield; despite the presence of O_2_. Pyruvate is converted to lactate, which requires all the glycolytic NADH output to be converted to NAD^+^, and this lactate is then excreted from the cell. So, aerobic glycolysis does not *net* output any NADH.

## Cancer cells use aerobic glycolysis some or all of the time

Cancer cells produce ATP through aerobic glycolysis (Warburg effect) some or all of the time [3-7]. This metabolic fingerprint is already exploited clinically to diagnose cancer using the radiolabeled glucose analogue - ^18^Fluoro-deoxyglucose (FDG) - in positron emission tomography (PET) [8]. Cancer cells utilising aerobic glycolysis have higher glucose uptake rates than surrounding, untransformed cells utilising aerobic respiration. It is this disparity that permits tumour identification. The vast majority of metastatic tumours are highly glycolytic (>90%) [9-10] and it may be that all of them are, but not all can be resolved at the present sensitivity and selectivity of the technique [11] e.g. those smaller than 0.8 cm^3^.

Warburg suggested that cancer cells use aerobic glycolysis because their OXPHOS apparatus is injured, and that this damage is a necessary step on the progression to a metastatic phenotype [3]. This is not always true. Yes, some cancer cell lines use aerobic glycolysis always and some do have defective OXPHOS apparatus. However, others use it only for a proportion of the time; using aerobic respiration at other times [6, 12-14]. They can switch between glycolytic and oxidative metabolism in a reversible fashion. For example, the AS-30D rat hepatoma cell line uses aerobic glycolysis for 60% of the time and aerobic respiration for the remainder [15]. A tumour of this cell line should still always be identifiable by PET because the cancerous cells are not all in metabolic sync – at any one time, a proportion of the cells will always be expressing aerobic glycolysis even if others are utilising aerobic respiration. Some cancer cells can utilise OXPHOS but exhibit a glucose-induced switch into aerobic glycolysis (Crabtree effect [16]).

## Why do cancer cells employ aerobic glycolysis?

Why do cancer cells utilise aerobic glycolysis some or all of the time? I propose that they need to use aerobic glycolysis, at the very minimum, during some crucial points in their proliferation cycle; for example, during DNA replication (S phase). This is a deduction from the fact that aerobic glycolysis is not restricted to cancer cells, but is applied by other highly proliferating cells e.g. embryonic cells [17-18]. For example, proliferating thymocytes switch from oxidative to glycolytic metabolism [19-24] during the G_1_/S transition of their cell cycle [19]. Their increasing dependence upon glycolysis peaks in concert with DNA replication [20]. OXPHOS produces harmful Reactive Oxygen Species (ROS), in addition to useful ATP, and these can damage cellular componentry. I propose that this is a tolerated evil normally but when cells are proliferating at an appreciable rate, and significant DNA replication is occurring, cells deliberately shunt OXPHOS. They do this by switching into aerobic glycolysis, which doesn’t produce or result in any NADH or FADH_2_ substrate for OXPHOS. In support, proliferating thymocytes employ aerobic glycolysis and produce minimal ROS in contrast to resting thymocytes, which use OXPHOS [24]. DNA replication is a vulnerable stage because DNA is unwound and exposed (from protective chromatin/histones [1-2]); in addition any damage at this stage could find itself reliably propagated into daughter cells, and further progeny downstream, before it can be identified as damage and repaired.

This peculiar metabolism may have an additional, insidious benefit for cancer cells. ROS are pivotal to apoptotic pathways [1, 2]. So with little or none produced, the cancer cell may then be liberated from the apoptotic contingency and free to conduct itself pathologically. Furthermore, although apoptosis is destructive, it is an active process and requires energy expenditure. By shutting mitochondria out of ATP generation it delimits their energy supply and may constrain their ability to initiate and pursue apoptosis. Furthermore, during aerobic glycolysis, the glycolytic enzyme hexokinase binds the voltage-dependent anion channel (VDAC) in the mitochondrial membrane and this suppresses apoptosis [25-26].

Aerobic glycolysis may also permit cancer cells to thrive in the hypoxia of solid tumours [27]. Furthermore, although it conveys a lower ATP yield than OXPHOS, it can produce ATP much faster (*100) [28] and so can in fact deliver more ATP than OXPHOS, if the substrate supply isn’t limiting. Cancer cells fast use of substrate denies it to surrounding untransformed cells and conveys a selective advantage [28]. Furthermore, it may produce local acidosis which may destroy adjacent, normal populations [11]. Greater glycolysis is good for proliferation because glycolytic intermediates are needed for macromolecular biosynthesis [17, 28]. They are needed for the pentose phosphate pathway that produces NADPH, which is needed for glutathione-dependent anti-oxidant mechanisms [29].

Cancer cells using aerobic glycolysis excrete lactate which may be then taken up and oxidised by other cancer cells using oxidative metabolism [30]. A given cancer cell may switch between the export and import of lactate, depending on where it is at in its proliferative cycle and whether it is employing aerobic glycolysis or oxidation. So, as a system, the tumour may in fact be gaining a full ATP yield from glucose. Incidentally, lactate transports (MCT 1 & 4) are an excellent drug target; although they will presumably require that the patient forgoes anaerobic exercise during treatment. An alarming heart rate monitor could be used to manage this constraint.

## If a cancer cell is switched out of aerobic glycolysis, into using OXPHOS, it can kill the cancer cell

As aforementioned, a cancer cell may use aerobic respiration for a significant proportion of its time. But I propose that at key times in the proliferating cycle it 

~~~
*mus*
~~~

t instead be using aerobic glycolysis and if it isn’t, it will be driven into apoptosis. There is experimental evidence for this.

Pyruvate is the end product of glycolysis. In aerobic glycolysis, pyruvate is converted to lactate and excreted. Lactate dehydrogenase (LDH) catalyses this reaction. In aerobic respiration, pyruvate dehydrogenase (PDH) out-competes LDH for pyruvate and switches it into the Krebs cycle. During aerobic glycolysis, PDH is inhibited by pyruvate dehydrogenase kinase (PDK), which can itself be inhibited by the dichloroacetate (DCA) drug [31]. So, the DCA drug can switch a cell from aerobic glycolysis to oxidative respiration, with its constituent Krebs and OXPHOS components. This drug induced conversion switches cancer cells into apoptosis [31]. Indeed, DCA in the drinking water of rats can prevent and reverse tumour growth 

~~~
*in vivo*
~~~

. DCA is an exciting cancer drug prospect and is already in clinical use for other applications. OXPHOS producing ROS is the trigger for apoptosis in these cancer cells [31].

Why should OXPHOS producing ROS be doomsday for cancer cells but permissible for non-transformed cells? I propose that it relates to proliferation. When cells are proliferating and DNA is unwound, unprotected and being replicated: if high ROS levels are sensed at this time, the process is aborted and the cell is driven into apoptosis. At other times, outside of this key stage(s) in the proliferation cycle, ROS are tolerated and the cell may utilise OXPHOS. The duration of this sensitive period may be quite narrow. I suggest that if one makes a proliferating cell constitutively use OXPHOS, robbing it of the ability to switch off OXPHOS at key stages, one will push it into apoptosis. Or at the very least, halt its proliferation. There is experimental evidence supporting the former [31] and the latter [23]. The latter study relates to healthy proliferating cells, rather than cancer cells. It shows that these issues of aerobic glycolysis are wider than cancer cells and may encompass all highly proliferating cells.

In conclusion, I propose that the use of aerobic glycolysis by cancer cells is not a function of their pathology but is a normal, physiological state employed by highly proliferating cells. If they are not in this state during key stages of the proliferating process, there are other physiological mechanisms that will halt their proliferation or drive them into apoptosis. This is a safety measure to ensure that ROS don’t damage DNA during the DNA replication process. DNA damage cannot be afforded at this stage because there isn’t the time for it to be repaired before the DNA strands are separated, and with this separation the record of damage is lost forever and the damage is rendered into permanence.

## Further evidence

Increasing the NAD^+^ level, by a treatment with its pre-cursers, is observed to have an anti-cancer effect [32]. The last step of aerobic glycolysis produces lactate and NAD^+^. I propose that with an imposed, increased NAD^+^ level, this reaction is more unfavourable. Especially as compared to the competing reaction: so pyruvate and NADH are more liable to enter the Krebs cycle and OXPHOS respectively. ROS are then produced that halt proliferation and trigger apoptosis. This rationale also explains why [32] found increased Complex I activity to be unconducive to cancer proliferation and survival.

Friedreich ataxia is an inherited neurodegenerative disorder caused by a reduced expression of frataxin, which results in less OXPHOS and ATP synthesis. Overexpressed frataxin stimulates OXPHOS, and ATP production, in colon cancer cells [33]. This stimulation suppresses colony formation 

~~~
*in vitro*
~~~

 and tumour formation/growth in nude mice; “these results support the view that an increase in oxidative metabolism (induced by mitochondrial frataxin) may inhibit cancer growth in mammals” [33].

## NADH as a cancer cure

I propose that exogenous NADH administration will cure some cancers. NADH is a substrate for OXPHOS and a continuous supply will confer a constitutive, enduring utilisation of OXPHOS in cancer cells. This will either halt their proliferation or trigger apoptosis.

Exogenous NADH will be harmless to most normal cells. It will just provide an additional source of energy for them. They will lower their glycolytic and TCA cycle rate accordingly to produce less NADH themselves (via ATP’s allosteric feedback mechanisms upon key glycolytic enzymes [1, 2]). Exogenous NADH may, however, be harmful to non-malignant, proliferating cells. So, there could be side effects. Although, in an adult, the proportion of healthy cells that proliferate at high rates is very small. So, these are likely to be slight.

The anti-cancer effect of a number of conventional treatments is based on their ability to stimulate ROS production and then apoptosis [34] e.g. radiation, etoposide, arsenates, vinblastine, cisplatin, mitomycin C, doxorubicin, camptothecin, inostamycin, neocarzinostatin etc. These typically stimulate ROS production, and damage, across all cells: cancerous or healthy. Cancer cells may be affected more because proliferating cells have a lower tolerance for ROS before apoptosis is initiated, as discussed. But normal cells are also damaged and the side effect profiles can be horrendous. Contrast this with the more targeted action of NADH therapy.

## NADH delivery

How can NADH be delivered into cells? NADH transports can be expressed at the surface of eukaryotes [35] and prokaryotes [36]; extracellular NADH can reach the OXPHOS apparatus and increase intracellular ATP [37]. NADH can be successfully administered orally or by intravenous/intraperitoneal infusion [38].

## NADH may combat the weight loss, wastage and weakness of cancer patients

Exogenous NADH may help the energy state of tumour bearing organisms, which typically suffer severe weight loss, wastage and weakness [39]. This is a seriously debilitating and upsetting aspect of cancer for patients. It can even be the cause of death. Wastage may happen because of the inefficiencies of cancer metabolism and the requirement to recycle glucose from lactate in the liver. This requires 6 ATP per lactate molecule (Cori cycle, [1-2]) whereas a molecular conversion of glucose to lactate in cancer cells only yields 2 ATP.

ATP infusions have been trialled to remedy this wastage of cancer patients, with some success [40-47]. However, it is unclear how ATP can enter cells. It is widely believed that mammalian cells cannot import external ATP (although this has been disputed, [47]). NADH therapy may produce better results as there is thought to be NADH transports at mammalian cellular surfaces [35, 37] and NADH can be given orally [38] rather than by long – and so by necessity, infrequent - periods of intravenous infusion.

## NADH will not work for cancer cell lines that have a defective OXPHOS apparatus

Some cancer cells have a working OXPHOS apparatus and some may use it some of the time. Others have a defective OXPHOS apparatus and are wholly reliant on aerobic glycolysis. These latter lines will be resistant to NADH treatment. However, their complete reliance on glycolysis will make them completely fallible to drugs that inhibit aerobic glycolysis [49-55] or glycolysis more generally. Although – note - drugs targeting generic glycolysis can cause severe side effects [56]. NADH therapy should be used first and if this fails then it can be surmised that the cancer cannot respire aerobically and is fully reliant upon glycolysis. Thus, its vulnerability is found. NADH therapy – with its (anticipated) low side effect profile - can be a curing step and, in failure, a screening step for lines susceptible to glycolytic inhibitors. Alternatively NADH therapy can be administered alongside a ketogenic diet (low glucose/carbohydrate, high fat), which may “starve” cells relying singly on glycolysis [57]. NADH, as a natural metabolite, may be permissible to communities and religions that prohibit the use of drugs, even in life threatening cases.

## FADH_2_ therapy

Exogenous FADH_2_ could be used as an alternative to NADH, which would circumnavigate Complex I and could be advantageous to destroying cancer cells that have developed a defective form of it. Although we would first need to discover whether exogenous FADH_2_ can passage the plasma and mitochondrial membranes.

## Prior work with NADH

NADH has been reported to have a kill effect upon some cancer cell lines 

~~~
*in vitro*
~~~

 [58] and in human case studies, where it was orally administered [59]. Unfortunately, the latter are anecdotal and lack rigor; further work is needed. The 

~~~
*in vitro*
~~~

 studies show, as anticipated, that NADH cannot kill all cancer cell lines. As aforementioned, I propose that those with a defective OXPHOS apparatus are immune to it.

## Conclusion

I propose that NADH pills will cure some cancers. A natural metabolite exerting such a specific kill effect is counterintuitive and intriguing. NADH therapy may also combat the wastage and weakness of cancer patients. Crucially, oral NADH has been proven as safe for humans in clinical trials (as a treatment for depression, Parkinson’s disease and chronic fatigue syndrome [60-63]).

This paper presents theory rather than data; but it is theory that is desperately needed at this point. Drug discovery should be driven by rationale rather than a random walk through compounds, awaiting fortuity and serendipity. We 

~~~
*must*
~~~

 rethink our approaches. We haven’t significantly improved the survival of Stage IV cancer patients since President Richard M. Nixon declared war on cancer over 40 years ago. My hope is that this article will prompt investigators to pursue directed animal and clinical studies.

